# *CotH* genes are necessary for normal spore formation and virulence in *Mucor lusitanicus*

**DOI:** 10.1101/2022.12.09.519853

**Authors:** Csilla Szebenyi, Yiyou Gu, Teclegiorgis Gebremariam, Sándor Kocsubé, Sándor Kiss-Vetráb, Olivér Jáger, Roland Patai, Krisztina Spisák, Rita Sinka, Ulrike Binder, Mónika Homa, Csaba Vágvölgyi, Ashraf S. Ibrahim, Gábor Nagy, Tamás Papp

**Author notes:** Corresponding author, (T.P.); or. These authors contributed equally to this work (Gábor Nagy, Tamás Papp).

## Abstract

Mucormycosis is an invasive fungal infection caused by certain members of the fungal order of Mucorales. The species most frequently identified as the etiological agents of mucormycosis belong to the genera *Rhizopus, Lichtheimia* and *Mucor*. The frequency of systemic mucormycosis has been increasing, mainly because of the elevating numbers of susceptible patients. Furthermore, Mucorales display intrinsic resistance to the majority of routinely used antifungal agents (e.g., echinocandins and short-tailed azoles), which limits the number of possible therapeutic options. All the above-mentioned issues urge the improvement of molecular identification methods and the discovery of new antifungal targets and strategies. Spore coat proteins (CotH) constitute a kinase family present in many pathogenic bacteria and fungi and participate in the spore formation in these organisms. Moreover, some of them can act as virulence factors being receptors of the human GRP78 protein during *Rhizopus delemar*-induced mucormycosis. We identified 17 *cotH-like* genes in the *Mucor lusitanicus* genome database. Successful disruption of five *cotH* genes was performed using the CRISPR-Cas9 system in *Mucor*. The CotH3 and CotH4 proteins play a role in adaptation to different temperatures as well as in developing the cell wall structure. We also show CotH4 protein is involved in spore wall formation by affecting the total chitin content and thus, the composition of the spore wall. Furthermore, we demonstrate that spore size is dependent on the *cotH4* gene. The role of CotH3 and CotH4 proteins in virulence was confirmed in two invertebrate models and DKA mouse model.

**IMPORTANCE:** Current treatment options for mucormycosis are inadequate resulting in high mortality rates especially among immunosuppressed patients. The development of novel therapies for mucormycosis has been hampered by lack of understanding of the pathogenetic mechanisms. The importance of the cell surface CotH proteins in the pathogenesis of *Rhizopus*-mediated mucormycosis has been recently described. However, the contribution of this family of proteins to the virulence of other mucoralean fungi and their functionality in vital processes remain undefined. Through the use of CRISPR-Case9 gene disruption system, we demonstrate the importance of several of the CotH proteins to the virulence of *Mucor lusitanicus* by using three infection models. We also report on the importance of one of these proteins, CotH4, to the spore wall formation through affecting the chitin content. Therefore, our studies extend the importance of CotH proteins to *Mucor* and identifies the mechanism by which one of the CotH proteins contributes to the development of a normal fungal cell wall, thereby indicating that this family of proteins can be targeted for future development of novel therapeutic strategies of mucormycosis.

---

Mucormycosis is a life-threatening opportunistic fungal infection caused by several members of the order Mucorales (1). *Rhizopus*, *Lichtheimia* and *Mucor* species have most often been isolated from such infections as the causative agents (2–6). These invasive infections, which can manifest as rhino-orbito-cerebral, pulmonary, gastrointestinal, cutaneous or disseminated diseases, are known for their aggressive progression and high mortality rates (i.e., 30-90% depending on the manifestation, the underlying condition of the patient and the therapy) (5, 7, 8). They most frequently occur in patients with an immunocompromised status due to immunosuppression (i.e., primarily for solid organ or hematopoietic stem cell transplantation) or hematological malignancies. Uncontrolled diabetes (with or without ketoacidosis), elevated levels of free iron in the blood and severe trauma can also be risk factors for mucormycosis (6, 7, 9, 10). The frequency of systemic mucormycosis has been increasing, mainly because of the elevating numbers of susceptible populations. Furthermore, Mucorales fungi display intrinsic resistance to the majority of routinely used antifungal agents (e.g., echinocandins and short-tailed azoles), which also limits the number of possible therapeutic options (4, 11). Recently, an increasing number of mucormycosis cases have been reported among COVID-19 patients treated with corticosteroids and with underlying diabetes (12–14).

Spore coat protein H (CotH) was first discovered in the endospore-forming bacterium *Bacillus subtilis* where it participates in the formation of the endospore coat (15). It proved to be an atypical protein kinase, which has an essential role in endospore formation by phosphorylating other structural proteins, such as CotB and CotG. Knock-out of the encoding *cotH* gene had a pleiotropic effect on the structure of the outer spore coat, as well as the development of a germination deficient phenotype (16–18).

CotH proteins occur not only in endospore-forming bacteria but are also present in many Mucoromycota species, such as *Mucor lusitanicus, Lichtheimia corymbifera, Cunninghamella bertholletiae, Rhizopus delemar, Saksanaea vasiformis, Syncephalastrum monosporum*, *Mortierella alpina* and *Umbelopsis isabellina* (19, 20). In the mucormycosis-causing *R. delemar* (synonym *R. oryzae*), several CotH proteins including, CotH1 (RO3G_05018; [EIE80313.1]), CotH2 (RO3G_08029; [EIE83324.1]) CotH3 (RO3G_11882; [EIE87171.1]), CotH4 (RO3G_09277; [EIE84567.1]); CotH5 (RO3G_01139); [EIE76435.1]); CotH6 (IGS-990-880_03186), CotH7 (IGS-990-880_09445), and CotH8 (IGS-990-880_11474) were previously identified (20, 21). Among them, CotH2 and CotH3 were found to mediate the interaction of the fungus with the glucose-regulated protein 78 (GRP78) expressed on the surface of endothelial cells (19), while CotH7 was found to mediate invasion of alveolar epithelial cells via interacting with integrin α3β1 (22). This interaction proved to be crucial for the fungal invasion of the host. Specifically, the level of GRP78 molecules significantly increases in sinuses, and lungs during diabetic ketoacidosis causing vulnerability towards the fungal infection (23). Moreover, IgG antibodies produced against a peptide of *Rhizopus* CotH3 protein and conserved among CotH7 protein protected mice with diabetic ketoacidosis (DKA) and neutropenia from mucormycosis (19). Thus, anti-CotH3 antibodies were proposed as promising candidates for immunotherapy treatment of human mucormycosis (1). Finally, due to the universal presence of *CotH* genes in Mucorales and the lack of their presence in other pathogens, they proved to be appropriate biomarkers for diagnosis through a PCR-based assay allowing fungal DNA detection in human urine samples (24).

*M. lusitanicus* (formerly *Mucor circinelloides* f. *lusitanicus* (25)) is a frequently used model organism in studying morphogenesis and pathogenesis of mucormycosis (26–31). In the genome of this fungus, we found 17 possible *cotH* genes, of which function and role in the pathogenicity or other mechanisms are yet unknown. This high number of genes lets us consider the possibility that the CotH family may be a diverse group of proteins having a role in various biological processes. The present study aimed to investigate the function and possible role in the pathogenicity of five *cotH* genes of *M. lusitanicus*.

## RESULTS

### *In silico* analysis of the CotH proteins

In the *M. lusitanicus* genome database (DoE Joint Genome Institute; *M. lusitanicus* CBS277.49v3.0; http://genome.jgi-psf.org/Mucci3/Mucci3.home.html), 17 potential CotH-like protein-coding genes were found by a similarity search using the amino acid sequence of *R. delemar* 99-880 CotH1 [EIE80313.1], CotH2 [EIE83324.1] and CotH3 [EIE87171.1], which were named as CotH1-CotH17, respectively (Table 1). *In silico* analysis of the *M. lusitanicus* putative CotH proteins amino acid sequences indicated that all of them carry the CotH kinase domain (Pfam: PF08757), while there are 15 genes in the *Rhizopus delemar* genome that carry the CotH domain (Pfam: PF08757) (15, 16). Furthermore, the *M. lusitanicus* CotH proteins are associated with the presence of a signal peptide, while only CotH2, CotH4, CotH10 and CotH12, contained sequences predicted to encode GPI-anchored proteins. Analysis of the sequences suggests that CotH proteins can be predominantly extracellular in nature. Among the 17 putative proteins, only the amino acid sequence of CotH4, CotH5 and CotH13 carry the special (M/Q/A-E/M/A-QTNDGAY-I/K-D-T/Y/G-N/A/E-E/N/T-N) motif previously described in *R. delemar* CotH2 and CotH3 proteins as a ligand for the GRP78 receptor (Fig. 1) (18), thereby suggesting that these proteins might play a role in the virulence of *M. lusitanicus*.

**FIG 1.**
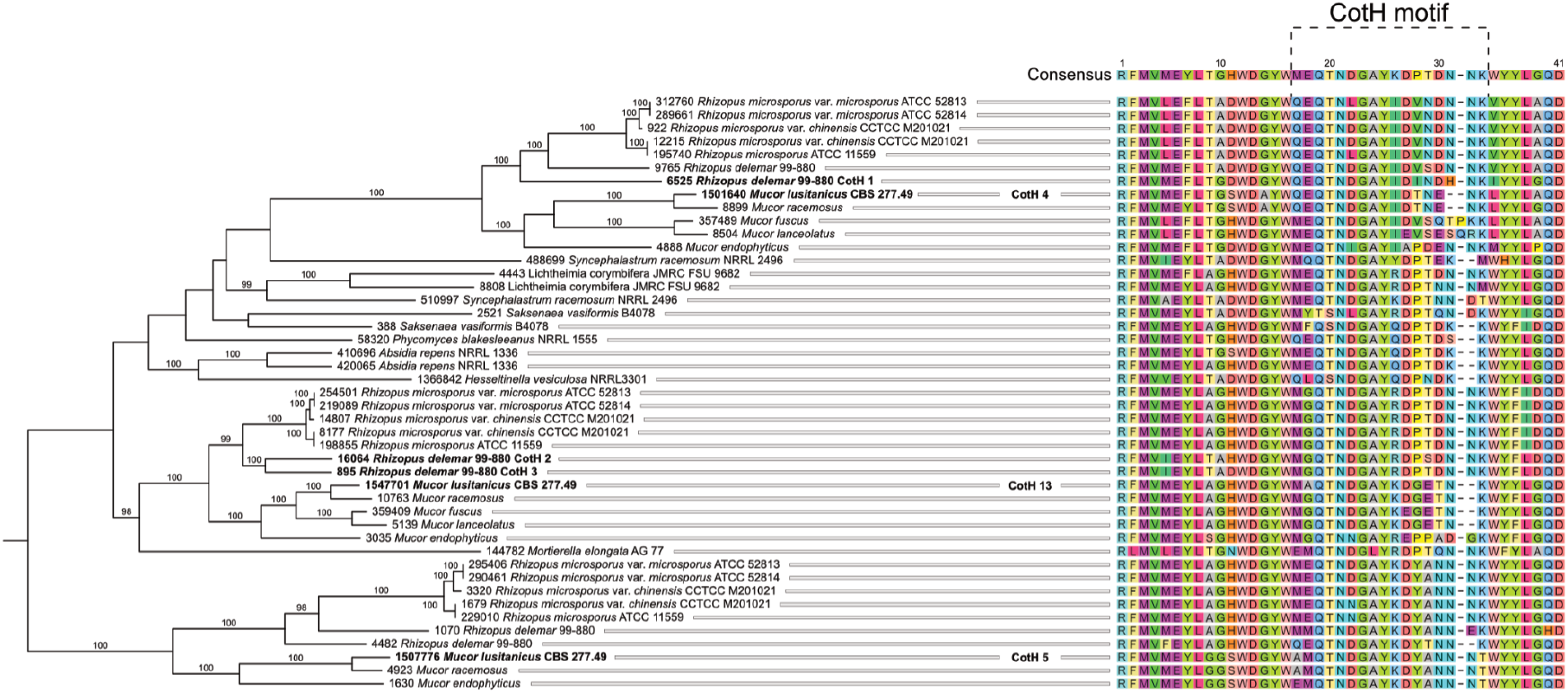
‘CotH-motif’ carrying proteins in Mucorales based on *in silico* analysis. The tree shows Clade 1 of the full phylogeny based on the available fungal genomic sequences presented in Fig. S1.

**TABLE 1.** The genomic position and protein ID in JGI MycoCosm’s database of the genes encoding the identified CotH-like proteins in *M. lusitanicus* and their names used in the study.

| Protein ID | Genomic location | Name in this study | Amino acid length (aa) | Signal sequence | GPI-anchor predicted |
| --- | --- | --- | --- | --- | --- |
| 1524506 | scaffold_5:3666732-3669040 | *cotH1* | 555 | + | - |
| 1443041 | scaffold_1:3072475-3074523 | *cotH2* | 621 | + | + |
| 109302 | scaffold_3:4206378-4208599 | *cotH3* | 707 | + | - |
| 1501640 | scaffold_4:354982-357045 | *cotH4* | 592 | + | + |
| 1507776 | scaffold_4:2485275-2486992 | *cotH5* | 524 | + |  |
| 1498659 | scaffold_3:4206283-4208688 | *cotH6* | 569 | + | - |
| 1461483 | scaffold_1:3050837-3053028 | *cotH7* | 594 | + | - |
| 1506605 | scaffold_4:2063119-2065181 | *cotH8* | 589 | + | - |
| 1590338 | scaffold_7:853960-855891 | *cotH9* | 595 | + | - |
| 1392021 | scaffold_7:437729-439702 | *cotH10* | 570 | + | + |
| 1395371 | scaffold_8:1812362-1814310 | *cotH11* | 596 | + | - |
| 1388805 | scaffold_5:3661652-3663532 | *cotH12* | 607 | + | + |
| 1547701 | scaffold_8:460886-463223 | *cotH13* | 590 | + | - |
| 1521260 | scaffold_5:2598993-2601087 | *cotH14* | 581 | + | - |
| 1407491 | scaffold_1:803076-804881 | *cotH15* | 581 | + | - |
| 1536541 | scaffold_6:3298508-3300483 | *cotH16* | 524 | + | - |
| 1561577 | scaffold_2:4842618-4846044 | *cotH17* | 590 | + | - |

All publicly available fungal genomic sequences were used for phylogenetic analysis of CotH proteins. The simplified view of the resulting tree is shown in Fig. 2 while the whole version can be found in Fig. S1. The collapse of the phylogenetic tree was based on the merging of clades of genes from predominantly closely related species. Interestingly, CotH proteins were found only in two phyla, Neocallimastigomycota and Mucoromycota and, within the latter, they were detected in two subphyla, Mucoromycotina and Mortierellomycotina. In the Mucoromycota linage, twelve well-supported clades with Mucorales CotH proteins occurred (Fig. 2). The remaining isolates, all of which were Neocallimastigomycetes, formed a distinct clade. Mortierellomycetes fungi are located in a self-clade [Clade 2], integrated between Clade 1 and Clade 3, formed by *Mucor* and *Rhizopus* isolates, respectively.

**FIG 2.**
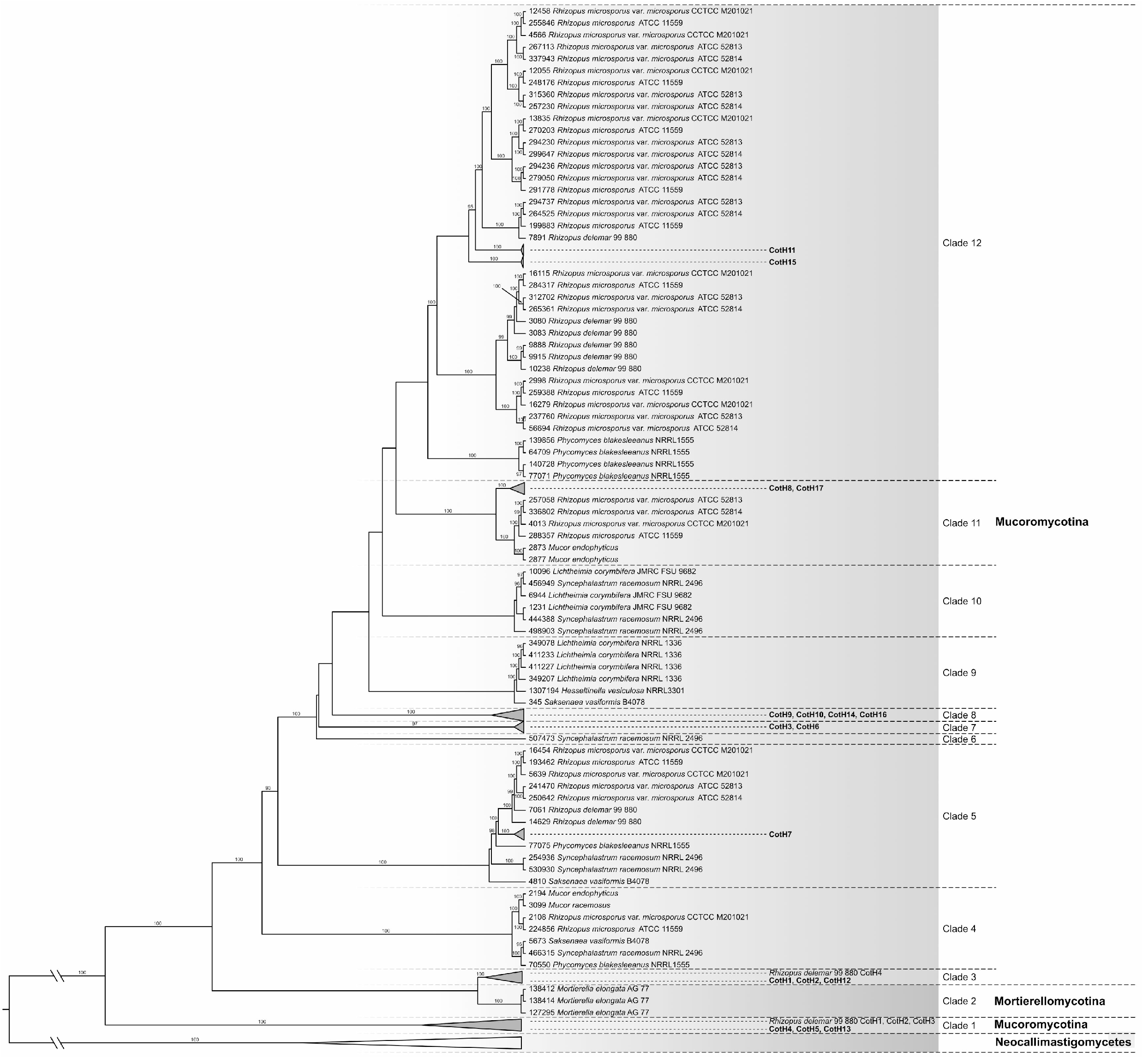
The collapsed version of Maximum Likelihood tree showing phylogenetic relationships of *cotH* gene sequences. The simplified view of the Maximum Likelihood tree was based on the merging of clades of genes predominantly closely related species. Only bootstrap values >95 % from maximum-likelihood analyses are shown above the branches.

*Mucor* CotH4, CotH5 and CotH13, which carry the special amino acid sequence described as a ligand for the GRP78 receptor (18) were localized in the same clade [Clade 1] together with the *Rhizopus* CotH2, and CotH3 proteins, which have been described previously for their essential role in virulence (Fig. 1) (18). Clade 1 also includes the *R. delemar* CotH7 ((IGS-990-880_09445), [protein ID: 9765]) that appears to be the major ligand mediating binding to integrin α3β1 of alveolar epithelial cells (22). Clade 3, Clade 4, Clade 5, Clade 11, and Clade 12 include both *Rhizopus* and *Mucor* CotH proteins, while some CotH proteins are segregated into clades formed exclusively by *Mucor* species [Clade 7 and Clade 8]. Uniquely, in Clade 1, some CotH homologues found in *M. lusitanicus* (i.e., CotH4, CotH5 and CotH13) are most closely related to the *R. delemar* and *P. blakesleeanus* orthologues than to their paralogs.

### Transcription analysis of the *M. lusitanicus cotH* genes

Transcription of the 17 *cotH* genes was analyzed by real-time quantitative reverse transcription PCR (qRT-PCR) on the second day of the cultivation at 28 °C (see Fig. S2 in the supplemental material), where *cotH2* and *cotH4* showed the highest expression. Most *cotH* genes reached their maximum transcript abundance on the second day of the cultivation at 28 °C, the optimum temperature of the fungus. Interestingly, transcription level of *cotH4* increased dramatically at higher glucose concentrations and in the presence of human serum (see Fig. S2 in the supplemental material). Based on the qRT-PCR analysis five genes (i.e., *cotH1* to *cotH5*) were selected for disruption and functional analysis.

### Knock-out of five *cotH* genes using the CRISPR-Cas9 system

Disruption of *cotH1*, *cotH2*, *cotH3*, *cotH4* and *cotH5* was performed using a CRISPR-Cas9 system and the resulting mutants were named as MS12-Δ*cotH1*+*pyrG*, MS12-Δ*cotH2*+*pyrG*, MS12-Δ*cotH3*+*pyrG*, MS12-Δ*cotH4*+*pyrG* and MS12-Δ*cotH5*+*pyrG*, respectively. For further analysis two independently derived mutants were selected. In each case, a template DNA containing the *pyrG* gene as a selection marker and two fragments homologous to the target site served as the deletion cassette. Transformation and genome editing frequencies are presented in Table S1 A. In the experiments performed to disrupt the *cotH5* gene, transformed protoplasts died before the colony development or colonies could be maintained only temporarily indicating that the disruption of this gene may be lethal for the fungus. It is also possible that the mutation causes a defect in protoplast regeneration, thus transformants could not be recovered even though the mutant would be viable under normal growth conditions. Investigating this possibility is hampered by the fact that currently only transformation of protoplasts of the fungus offers the possibility of producing stable genetic mutants of *M. lusitanicus*.

### Characterization of the knock-out mutants

#### Growth ability of the mutants

Growth ability of the mutants was examined at 20, 28 and 35 °C for four days. Growth analysis was performed with two independently derived mutants for each *cotH* knock-out. No significant differences in the growth of MS12-Δ*cotH1*+*pyrG*, MS12-Δ*cotH2*+*pyrG* and the control strain were observed at the optimum growth temperature of the fungus (28 °C) (Fig. 3A). However, the colony diameter of MS12-Δ*cotH3*+*pyrG* significantly increased, while that of MS12-Δ*cotH4*+*pyrG* significantly decreased compared to the control strain at this temperature (Fig. 3A). Apart from the growth intensity of these two strains, morphology of the mutants did not differ from that of the original strain. Cultivation at lower temperature did not affect the growth of the strains differently compared to the control (Fig. 3B). At 35 °C, growth of the *cotH4* disruption mutant was significantly less affected by the increased temperature than those of the *cotH1*, *cotH2* and *cotH3* mutants (Fig. 3C). Complementation of the *cotH3* or *cotH4* gene was achieved by an autosomally replicating plasmid construction, where the complementing gene construct was maintained extrachromosomally. After complementation of the *cotH3* and *cotH4* gene, the resulting MS12-Δ*cotH4*+*cotH4*+*leuA* strain showed the growth characteristics of the control (MS12+*pyrG*) (see Fig. 3).

**FIG 3.**
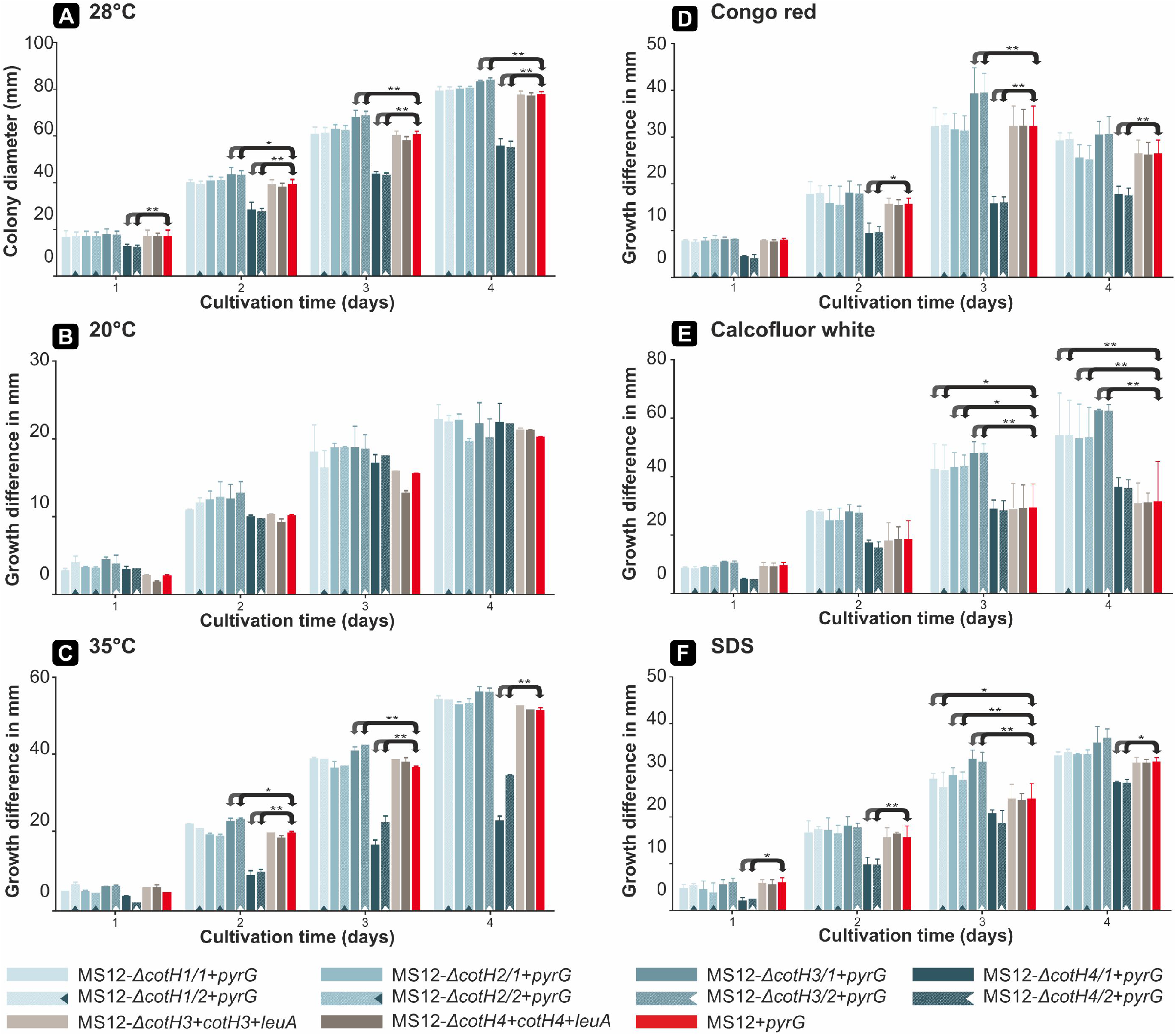
Growth of *cotH* mutants at different temperatures and effect of different stressors. (A) Colony diameter of the *cotH* mutant strains on leucine-supplemented minimal medium for 4 days at 28 °C. (B) Effect of the lower temperature on growth of *cotH* mutant strains on leucine-supplemented minimal medium for 4 days at 20 °C. (C) Effect of the higher temperature on growth of *cotH* mutant strains on leucine-supplemented minimal medium for 4 days at 35 °C. Effect of the lower (B) and higher (C) temperature on the growth of *cotH* mutant strains was represented by comparing the differences of the colony diameters measured after cultivation under the control conditions and those obtained in the cultivation under 20 °C and 35 °C. (D) Effect of Congo red (CR) on the growth of *cotH* mutant strains. (E) Effect of Calcofluor white (CFW) cell wall stressor on the growth of *cotH* mutant strains. (F) Effect of SDS membrane stressor on the growth of *cotH* mutant strains. The effect of the different stressors on the growth of the mutants was represented by comparing the differences of the colony diameters measured after cultivation under the control conditions and those obtained in the presence of the stressor. Values presented are from three independent cultivations (error bars indicate standard deviation). The strains were grown at 28 °C in the dark for 4 days, and colony diameters were measured daily. The effect of the different temperature (A-C) and stressors (D-F) was then plotted in millimeters and significance was calculated based on the effect of the temperatures or stressors on MS12+*pyrG*. *P* values calculated according to the statistical method two-sample, paired t-test. Values indicated with asterisks significantly differed from the value of the MS12+*pyrG* strain measured on the same day according to * *P* ≤ 0.05; ** *P* ≤ 0.01.

Cell wall stressors Congo Red (CR) and Calcofluor white (CFW) had significant effects on the growth of all strains compared to cultivation on normal medium (Fig. 3). Difference of the colony diameter of each mutant from that of the control strain on normal, untreated medium (Fig. 3A) (i.e., increased or decreased growth ability of the mutants) was taken into account during the evaluation. Using CR as a stressor, no difference in the growth intensity of the deletion mutants MS12-Δ*cotH1*+*pyrG* and MS12-Δ*cotH2*+*pyrG* was observed compared to MS12+*pyrG* (i.e., the control strain). MS12-Δ*cotH3*+*pyrG* displayed an increased sensitivity, while MS12-Δ*cotH4*+*pyrG* showed higher resistance to CR than the control strain (Fig. 3D). CFW exerted a significant effect on the growth of all *cotH* mutants. MS12-Δ*cotH1*+*pyrG*, MS12-Δ*cotH2*+*pyrG* and MS12-Δ*cotH3*+*pyrG* was more sensitive, while MS12-Δ*cotH4*+*pyrG* was more resistant to CFW than MS12+*pyrG* (Fig. 3E). The MS12-Δ*cotH4*+*pyrG* proved to be more resistant to SDS than the control strain. The effect of SDS was observed from the third cultivation day in the case of MS12-Δ*cotH1*+*pyrG*, MS12-Δ*cotH2*+*pyrG* and MS12-Δ*cotH3*+*pyrG* (Fig. 3F). For these strains, an increased sensitivity was observed. After complementation of the *cotH3* and *cotH4* gene, the resulting MS12-Δ*cotH3*+*cotH3*+*leuA* and MS12-Δ*cotH4*+*cotH4*+*leuA* strains showed the growth characteristics of the control (MS12+*pyrG*) (Fig. 3).

Under aerobic conditions, *M. lusitanicus* displays a hyphal growth while it grows in a yeast-like form under anaerobiosis. This feature, named as morphological dimorphism is regarded as an important property of pathogenicity (27). Gene disruption did not affect the anaerobic growth of any of the mutants compared to the control strain and yeast-like cells could be formed in the absence of the tested *cotH* genes. However, under microaerophilic conditions, MS12-Δ*cotH4*+*pyrG* produced more elongated hyphae than the control strain (see Fig. S3 in the supplemental material).

#### Transmission electron microscopic analysis of the spores of the mutant strains

To determine the effect of the gene disruptions on the spore wall structure of *M. lusitanicus* spores of the mutants and the control strain (MS12+*pyrG*) were subjected to transmission electron microscopic (TEM) image analysis (Fig. 4). Spore wall of MS12-Δ*cotH4*+*pyrG* showed a characteristic phenotype as it was abnormally thickened (Fig. 4B and D and File S1). After complementation of the *cotH4* gene disruption, the cell wall of MS12-Δ*cotH4*+*cotH4*+*leuA* spores was restored and the cell wall thickness of the complemented spores did not differ significantly from those of the control (MS12+*pyrG*) (see Fig. S4).

**FIG 4.**
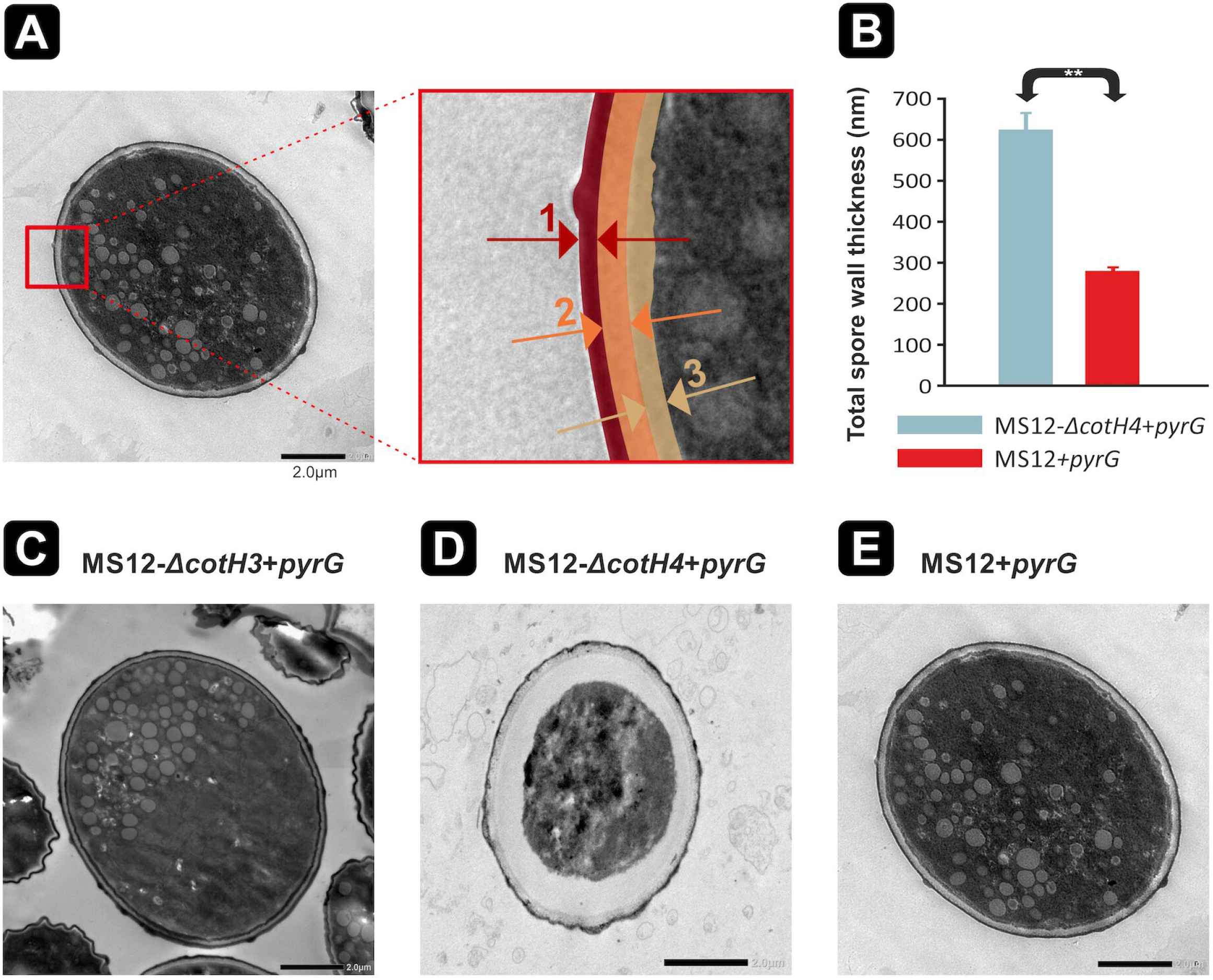
Cell wall changes of *cotH* mutants observed by TEM. (A) Schematic representation of the outer, inner, and middle layers of the cell wall of spores. TEM measurements represent the thickness of different cell wall layers. A/1 outer spore wall layer, A/2 middle spore wall layer, A/3 inner spore wall layer. (B) Total spore wall thickness of spores (1–3) compared to control strain. Spores of the *cotH* mutants (C) MS12-Δ*cotH3*+*pyrG*. (D) MS12-Δ*cotH4*+*pyrG*. (E) MS12+*pyrG* (control). Significance was calculated based on a two-sample, t-test, where ** *P* ≤ 0.001. Error bars indicate standard deviation. For each mutant, spore wall thickness was determined by measuring the thickness of five different spores at ten different points on the spore wall. Scale bars: 2 μm.

#### Surface analysis of fungal spores with fluorescent dyes

By monitoring the spores stained with CWF, we found that the dye-emitted intensity of the spores of MS12-Δ*cotH4*+*pyrG* significantly increased compared to the control (Fig. S5) suggesting a chitin accumulation in the spore wall. At the same time, staining for mannosyl derivatives using Concanavalin A-fluorescein isothiocyanate (ConA-FITC) did not reveal differences among the mutants and the control strain. Intensity of the *cotH4*-complemented (i.e., MS12-Δ*cotH4*+*cotH4*+*leuA*) spores did not differ significantly from those of the control (MS12+*pyrG*) (see Fig. S5 in the supplemental material).

#### *In vitro* interaction with macrophages

To examine whether recognition and internalization of the fungal spores by macrophages were affected by the disruption of the *cotH* genes in *M. lusitanicus*, the phagocytosing capacity (see Fig. S6 in the supplemental material) and the phagosome maturation of J774.2 cells exposed to the MS12+*pyrG* and the mutant strains were tested (see Fig. S7 in the supplemental material). Subsequently, J774.2 cells were co-incubated for three hours with the labelled spores and then the ratio of pHrodo^™^ Red + macrophages were examined by imaging flow cytometry (see Fig. S7A and B in the supplemental material). No significant differences were found between the mutant and the control strains in neither the phagocytic index values (see Fig. S6C in the supplemental material). Similarly, testing the survival of fungal spores co-cultured for 3 h with murine J774.2 cells did not detect differences between the control and the mutant strains (see Fig. S6D in the supplemental material). Moreover, our results suggested that the viability of spores did not decrease in the presence of macrophages at all. Furthermore, acidification of phagosomes is not affected by the lack of the examined CotH proteins (see Fig. S7B in the supplemental material) and the absence of CotH proteins did not affect the survival of spores after *in vitro* interaction with macrophages.

#### *In vivo* study of virulence of *cotH* mutant strains

To investigate the role of CotH proteins in pathogenicity of *M. lusitanicus*, *Drosophila melanogaster* and wax moth larvae (*Galleria mellonella*) were used as non-vertebrate animal models in *in vivo* virulence studies.

In *D. melanogaster* (Fig. 5A), pathogenicity of the control strain MS12+*pyrG* and the wild type CBS277.49 did not differ significantly. However, lack of the *cotH3* and *cotH4* genes significantly decreased the virulence in the mutants compared to the controls. In case of the *Galleria* model Fig. 5B, disruption of the *cotH4* gene resulted in significantly decreased virulence of the mutant strain. In an intratracheally infected DKA mouse model, all the mice infected with the wild type *M. lusitanicus* strain CBS277.4 died before the third-day postinoculation (Fig. 5C). In this model, MS12-Δ*cotH3*+*pyrG* and MS12-Δ*cotH4*+*pyrG* showed significantly decreased virulence. After complementation of the *cotH3* and *cotH4* gene, the virulence of the resulting MS12-Δ*cotH3*+*cotH3*+*leuA* and MS12-Δ*cotH4*+*cotH4*+*leuA* strains showed the characteristics of the control (CBS277.49) (Fig. 5).

**FIG 5.**
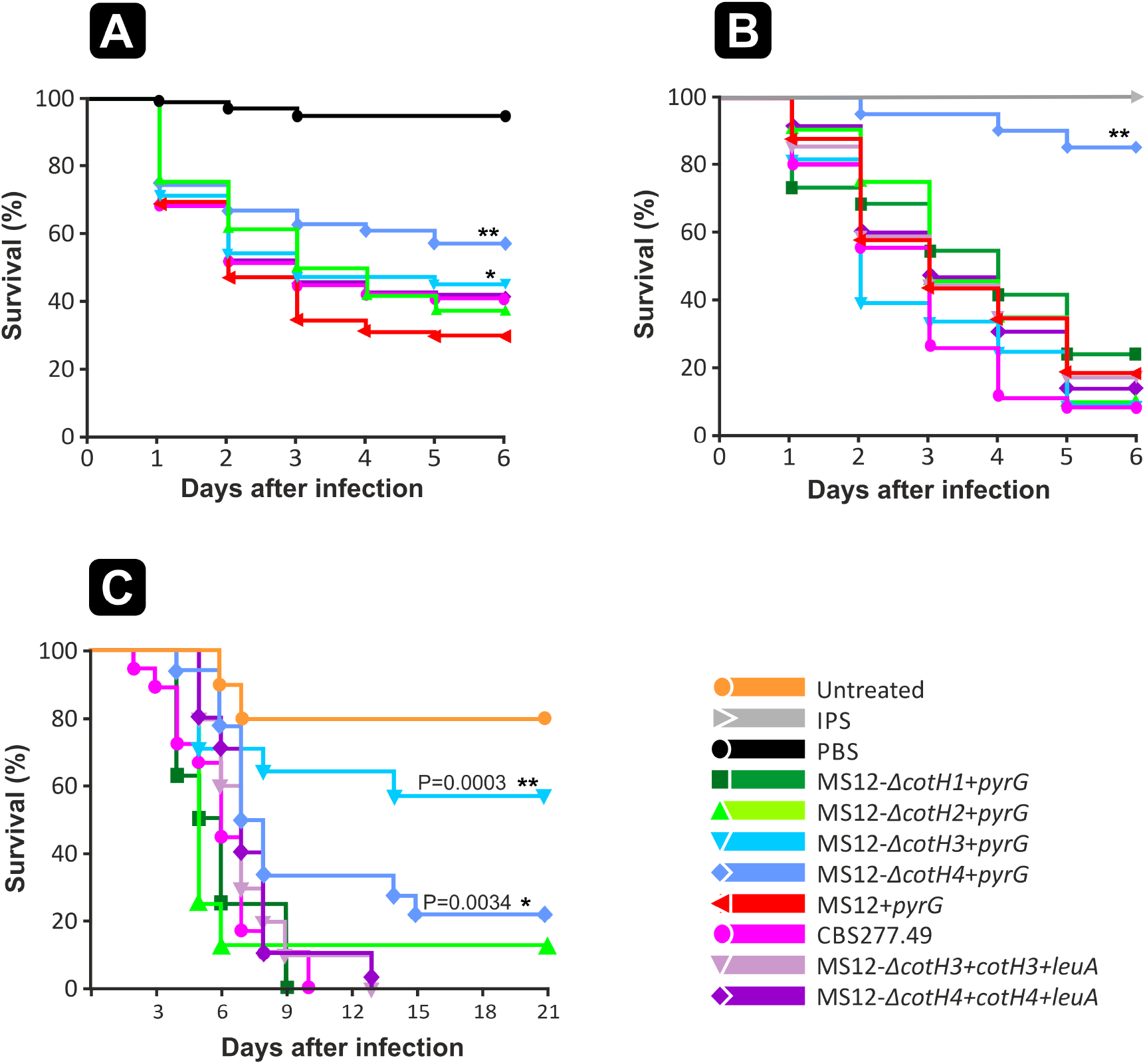
Virulence studies with *cotH* mutants. (A) Survival of *Drosophila melanogaster* (n=60) infected with the *cotH* mutants and the control *M. lusitanicus* MS12+*pyrG* and CBS277.49 strains. Survival curve followed by asterisks were significantly differed from the control strain (MS12+*pyrG*) according to the Log-rank (Mantel-Cox) test (* *P* ≤ 0.05 and ** *P* ≤ 0.001). The results summarize the results of 3 independent experiments. (B) Survival of *Galleria mellonella* (n=20) infected with the *cotH* mutants and the control *M. lusitanicus* MS12+*pyrG* and CBS277.49 strains. Survival curve followed by asterisks were significantly differed from the control strain according to the Log-rank (Mantel-Cox) test (** *P* ≤ 0.001). The results summarize the results of 3 independent experiments. (C) Virulence of *cotH* mutants in a DKA mouse model following intratracheal infection. Male DKA Male ICR (CD-1®) outbred mice (≥20 g) (Envigo) (n=8) were infected intratracheally with 2.5×10^6^ fresh spores (1X10^8^ spores / ml) in 25 μl PBS. The results summarize the results of 2 independent experiments. Survival curve followed by asterisks were significantly differed from the control strain according to the Log-rank (Mantel-Cox) test (* *P* ≤ 0.05; ** *P* ≤ 0.01).

## DISCUSSION

CotH protein encoding genes are widely present in Mucorales fungi (19). Although present in other organisms, Mucorales *cotH* genes have diverged sequences from their orthologs in *Bacillus* species. However, the function of *cotH* genes has been verified in only a few of them (e.g., *B. subtilis, B. cereus* and *R. delemar*) (15, 16, 19, 32). For instance, the *Rhizopus* CotH3 carries the amino acid sequence “MGQTNDGAYRDPTDNN” which is assumed to be a key factor in the specific interaction between the fungal cells and the host’s endothelial cells via the GRP78 molecule (19). This interaction was shown to be critical in enhancing *R. delemar* virulence through promotion of haematogenous dissemination (18). Equally important, the ability of *R. delemar* spores to invade and damage nasal epithelial cells was directly proportional to the expression of GRP78 on nasal epithelial cells triggered by host conditions mimicking hyperglycaemia and ketoacidosis. Thus, explaining the increased susceptibility of diabetics in ketoacidosis to the rhino-orbito-cerebral form of mucormycosis (ROCM) (22). Concordant with these findings, is the increased number of mucormycosis cases (mainly ROCM) detected among COVID-19 patients treated with corticosteroids and with underlining diabetes (i.e. COVID-19-associated mucormycosis [CAM]) (33). Interestingly, significantly higher serum GRP78 levels in COVID-19 patients have been reported and could explain the increased incidents of CAM through CotH/GRP78 mediated invasion of host tissues (34). Thus, some members of the CotH protein family serving as ligands for GRP78 receptors may be considered as potential therapeutic targets (1).

In a previous study, several different *cotH* transcripts were reported to be expressed in *M. lusitanicus* using the domain profile PF08757 (34). Moreover, it was also noted that the number of *cotH* transcripts was two times higher in the transcriptome of *Mucor* strains considered to be pathogenic, than in *Mucor* strains used in cheese production (35). This study used “CotH motif” described previously for *Rhizopus* (35) to report on the presence of three CotH-proteins in *M. lusitanicus*. In our study, we found 17 genes in *M. lusitanicus*, which contain the PF08757 domain, of which three genes (i.e. *cotH4, cotH5* and *cotH13*) harbor the so-called “CotH-motif”. In the phylogeny of the *cotH* genes, *M. lusitanicus cotH4, cotH5* and *cotH13* localized in the same clade [Clade 1] together with the *R. delemar cotH2* and *cotH3*. Only the genes positioned into the Clade 1 of this phylogeny contained the CotH-motif, suggesting that these genes may have a role in the CotH–GRP78 interaction.

Expression of the *M. lusitanicus* CotH3 protein proved to be essential for the virulence in DKA mouse model. However, the encoding gene localizes to Clade 7 along with other CotH proteins and does not carry the CotH-motif predicted to interact with GRP78. This result suggests that host targets other than GRP78 are involved in mucormycosis pathogenesis due to *M. lusitanicus*. Indeed, integrin α3β1 have been reported to be the target for *R. delemar* CotH7, which results in activation of the epidermal growth factor receptor (22, 36).

Phylogenetic analysis indicated that CotH proteins show a strong dominance in the Mucoromycotina fungi within the fungal kingdom. Clade 4, Clade 5, Clade 11, and Clade 12 include both *Rhizopus* and *Mucor cotH* genes, while some *cotH* genes are segregated into clades formed exclusively by *Mucor* species [Clade 7 and Clade 8] suggesting various gene duplication events within the Mucoromycota clade. Distribution of the genes in the phylogeny also suggests that several duplication event may occur in the zygomycete ancestor before the divergence of species. The presence of this large number of genes and their phylogenetic analysis led us to consider the possibility that the CotH family is a diverse group of proteins, which may be involved in many biological processes and not exclusively in the virulence.

Since orthologs of CotH proteins are mostly considered as spore surface or spore envelope structural elements (35), the spore wall composition of the fungus and its mutants generated in this study was monitored. TEM images revealed the separation of the spore wall and the cell membrane of the *cotH4* disruption mutant. Analysis using fluorescent dyes specifically binding to the cell wall components (37, 38) indicated a chitin accumulation between the two structures. Changes in the cell wall composition can alter the susceptibility to cell wall stressors, such as CFW and CR (37) and may influence the fungal virulence (39). Indeed, disruption of the *cotH4* gene decreased the sensitivity to CR and caused resistance to other stressors (i.e., CFW and SDS) at the same time, MS12-Δ*cotH3*+*pyrG* displayed an increased sensitivity to CR. The ability of pathogenic fungi to respond to stress is crucial in adapting to the host environment (40). Our results suggest that the proteins encoded by the *cotH3* and *cotH4* genes are involved in the determination of the spore wall composition and structure.

Chemical components, such as the surface molecules, and physical properties, such as the size and shape, of the fungal propagules greatly influence pathogen recognition and further removal of the fungus by the immune cells (41). Changes in the spore wall of the *cotH4* disruption mutant, as well as the absence of the CotH proteins did not affect their recognition by the macrophages, also, no significant difference was found in the number of fungal cells phagocytosed by a macrophage. The ability of the spores to germinate and penetrate the cells determines the further outcome of the infection (41). It was previously described that *Mucor* spores within the phagosome are exposed to a cytotoxic environment, which includes acidification, nutrient starvation, oxidation, and the presence of antimicrobial proteins (41). We examined whether the acidification of phagosomes could be affected by the CotH proteins. In our experiments, macrophages were equally capable to kill the spores of the *cotH* knock-out mutants, so neither the recognition nor the killing mechanism was impaired or altered. Despite the involvement of CotH proteins in cell wall remodelling and/or integrity, there is no indication that CotH1-4 proteins would be involved in phagosome maturation blockade in *Mucor*, as suggested by a previous study (41). It should be noted, however, that of the 17 CotH proteins encoded in the *Mucor* genome, two other proteins (i.e., ID: 76509, CotH8 and ID: 166651, CotH17) are suggested to play a role in macrophage interactions (41).

Although the information on the role of genes or proteins involved in infection can be obtained through *in vitro* experiments, the function of systemic infection can only be elucidated in an *in vivo* model (42). Disruption of both the *cotH3* and the *cotH4* genes caused reduced virulence in the *Drosophila* infection model, while the disruption of *cotH1* and *cotH2* genes did not affect the virulence of the fungus. The *Drosophila* pattern recognition receptors after the recognition of conserved microbial patterns can activate a cellular and humoral response that is specific to a particular microorganism (43). The wax moth, *G. mellonella* is also a widely used non-vertebrate model organism to examine the pathogenicity of different filamentous fungi such as Mucorales species (26, 31, 44). *In vivo* viability studies in *G. mellonella* also confirmed the role of CotH4 protein in virulence. The pathological alterations that resemble human mucormycosis are limited if we would like to use *Drosophila* or *Galleria*, but an appropriate model for testing larger numbers of mutants (42). Furthermore, there is a limitation of the use of invertebrate hosts is whether the results obtained in these models can be adapted to mammals and the human body because of fundamental differences such as the lack of an adaptive immune system and specific organs (42).

Inhalation of Mucorales spores is the most common route of infection. Thus, intratracheal instillation or intranasal inhalation is the most widely used models to mimic pulmonary infection. Disseminated mucormycosis can also be induced by intravenous inoculation of the fungus, usually into the tail vein and mimicking direct inoculation into the blood as the infection occurs in severe trauma (45). Therefore, the role of *cotH* genes in *R. delemar* was investigated using a DKA mouse model in which pulmonary mucormycosis was induced by intratracheal instillation of fungal spores (19). This model is preferably used to investigate the pathomechanism of mucormycosis, as the most common underlying condition of this fungal disease is diabetic ketoacidosis besides immunodeficiency (3,30,38). For *M. lusitanicus*, an intratracheal mouse infection model has not been previously used, so the infectivity of WT (CBS277.4) in mice was first established (31). Viability studies in DKA mice demonstrated that deletion of either CotH3 or CotH4 genes attenuate the pathogenicity of *M. lusitanicus*. Importantly, the *cotH3* mutant did not show reduced virulence in a *Galleria mellonella* model but did in a DKA mouse model with elevated GRP78 receptor expression, although the protein does not carry the characteristic motif determined earlier for *R. delemar*. The data suggest that *cotH* mutants exhibit altered cell wall composition or organization, with the *cotH4* mutant showing a significant loss of cell wall integrity and cell wall chitin composition. Host recognition of the fungal cell wall often determines the outcome in the host and plays an important role in the pathomechanism of the infection. Based on these findings, it seems conceivable that in *M. lusitanicus*, CotH3 and CotH4 proteins mediate the process of fungal infection in a cell wall-dependent manner.

Spore size dimorphism is linked to virulence of *M. lusitanicus* species (24). However, considering that the mutant lacking the CotH3 protein showed reduced virulence in the pathogenicity studies, but no change in spore size, it can be assumed that spore size is not the only determining factor of virulence. Interestingly, interaction with the GRP78 receptor is unlikely in case of this mutant, as the CotH3 protein does not carry the “CotH-motif”. Alternatively, other motifs that are yet to be identified bind to GRP78. Due to its sequence similarity to the *Rhizopus* CotH3 and the presence of the “CotH-motif”, CotH4 could be a potential ligand for the GRP78 receptor, which requires further investigation. Given that CotH proteins are involved not only in pathogenicity but also in spore structure, it is important to consider the role of CotH protein family members not only as virulence factors but also in spore formation and other physiological roles.

## MATERIALS AND METHODS

### Strains, media and growth conditions

The strain MS12 of *Mucor lusitanicus* (formerly known as *Mucor circinelloides* f. *lusitanicus*) is auxotrophic to leucine and uracil (*leuA*^-^ and *pyrG*^-^) and derived from the strain CBS277.49 by chemical mutagenesis (46), was used in the transformation experiments. As the lack of a functional *pyrG* slightly affects the growth and virulence of *M. lusitanicus* (47), the strain MS12+*pyrG*, was used as a control during the characterization of mutants. In this strain, uracil auxotrophy was complemented by expressing the *pyrG* gene (47). In certain experiments, CBS277.49 was also involved as a control. Growth analysis was performed with two independently derived mutants for each *cotH* mutations.

For qPCR experiments, human serum was isolated from venous blood of the same donors taken into serum separation blood collection tubes (BD Vacutainer, Becton Dickinson, Franklin Lakes, NJ, USA). Then, coagulation tubes were centrifuged at 300 g for 15 min at room temperature and the serum was collected and added in 10%, then cultivation was performed for two days at 28 °C in liquid minimal media.

For nucleic acid extractions, spores were plated onto solid minimal medium (YNB; 10 g/l glucose, 0.5 g/l yeast nitrogen base without amino acids (BD Difco, Becton Dickinson, Franklin Lakes, NJ, USA), 1.5 g/l (NH_4_)_2_SO_4_, 1.5 g/l sodium glutamate and 20 g/l agar) supplemented with leucine and/or uracil (0.5 mg/ml), if required and incubated at 28 °C for 4 days. To test the mitotic stability of the transformants, malt extract agar (MEA; 10 g/l glucose, 5 g/l yeast extract, 10 g/l malt extract and 20 g/l agar) was used as a complete, non-selective medium. To examine the effect of the temperature on the growth, 10^4^ spores were plated on solid YNB and incubated at 20, 28 and 35 °C. Anaerobic growth was performed in a BBL GasPak Anaerobic System (Becton Dickinson) with Anaerocult A (Merck, Darmstadt, Germany) at 28 °C. Microaerophile growth was performed in a BBL GasPak Anaerobic System (Becton Dickinson) with Anaerocult C (Merck, Darmstadt, Germany) at 28 °C, where 10^4^ spores were plated on solid YNB and incubated at 28 °C for two days. Fungi grown on YNB under anaerobiosis were sampled on the second day of culture and the morphology of the fungal cells was examined by light microscopy. To determine the effect of membrane and cell wall stressors, 10^4^ spores were point-inoculated at the centre of YNB with or without the stressor, which were sodium dodecyl sulfate (SDS; 4 mg/ml), Congo red (CR; 2 mg/ml) and Calcofluor white (CFW; 0.1 mg/ml). For growth tests, plates were incubated at 28 °C for 4 days, in dark and colony diameter was measured daily. In each case, the difference in the colony diameters of the mutant strains and that of the control was determined under the control conditions (i.e., when the strains were grown at 28 °C on YNB) and in the presence of the stressor. The effect of the different temperatures and stressors on the growth of the mutants was represented by comparing the differences in the colony diameters measured after cultivation under the control conditions and those obtained in the cultivation under stressors. The effect of the different temperatures was then plotted in millimeters and significance was calculated based on the cultivation under different temperatures on MS12+*pyrG*.

### Sequence and phylogenetic analysis of CotH proteins

Motifs, domains and main features of the CotH proteins were predicted using the tools available at the Expasy Bioinformatics Resource Portal (http://www.expasy.ch), such as Compute pI/Mw, MyHits, PROSITE and ProtScale (48). For the phylogenetic analysis, BLASTp search was conducted on the JGI MycoCosm portal (https://mycocosm.jgi.doe.gov/mycocosm/home)(49) with 17 sequences of *Mucor lusitanicus* CBS277.49 containing CotH domain. All retrieved sequences were scanned for CotH domains with InterProScan 5.48-83.0 (50) based on Pfam database (51). Only those sequences were retained, which contained CotH domains solely. Filtered sequences were clustered by using MMseq2 v. bbd564172bd55d9e6acd1170e59790c37157a21b (52). with default settings. Multiple sequence alignment was conducted using MAFFT v. 7.453 (53) with the E-INS-i iterative refinement method including sequences from all clusters which contained CotH domain based on the clustering results. Phylogenetic reconstruction was carried out by using IQ-TREE v. 1.6.12 (54) with the LG4M+R7 model determined by the inbuilt ModelFinder tool (55). Statistical support of the best tree was calculated with ultrafast bootstrap approximation (56) in 5000 replicates.

### General molecular techniques

Genomic DNA and total RNA were isolated using the ZR Fungal/Bacterial DNA MiniPrep (Zymo Research, Irvine, CA, USA) and the Direct-zol RNA MiniPrep (Zymo Research, Irvine, CA, USA) kits, respectively, according to the instructions of the manufacturer. To amplify genes or gene fragments from genomic DNA, the Phusion High Fidelity DNA Polymerase (Thermo Fischer Scientific, Waltham, MA, USA) was used according to the manufacturer’s recommendations. PCR products were isolated and concentrated using the Zymoclean Large Fragment DNS Recovery Kit (Zymo Research, Irvine, CA, USA) and DNA Clean & Concentrator-5 (Zymo Research, Irvine, CA, USA). Restriction digestions and ligations were carried out according to the commonly used methods (57). To clone the PCR fragments, the pJET1.2/blunt vector (CloneJET PCR Cloning Kit, Thermo Fischer Scientific, Waltham, MA, USA) was used according to the manufacturer’s instructions. Plasmid purification was carried out using the GeneJET Plasmid Miniprep Kit (Thermo Fisher Scientific, Waltham, MA, USA) according to the manufacturer’s recommendations. Sequencing of the cloned fragments was commercially performed by the LGC Genomics (Berlin, Germany). Sequences obtained were aligned by using the BioEdit 7.2 sequence editor program (58) and analyzed using the Basic Local Alignment Search Tool (BLAST) at the site of the National Center for Biotechnology Information (NCBI) (https://blast.ncbi.nlm.nih.gov/Blast.cgi). Oligonucleotide sequences were designed based on the sequence data available in the *M. lusitanicus* CBS277.49v3.0 genome database (DoE Joint Genome Institute; http://genome.jgi-psf.org/Mucci2/Mucci3.home.html) (59). Primers used in the study are listed in Table S1B.

### qRT-PCR analysis

Reverse transcription was carried out with the Maxima H Minus First Strand cDNA Synthesis Kit (Thermo Fischer, Waltham, MA, USA) using random hexamer and oligo(dT)_18_ primers, according to the manufacturer’s recommendations. qRT-PCR experiments were performed in a CFX96 real-time PCR detection system (Bio-Rad) using the Maxima SYBR Green qPCR Master Mix (Thermo Fischer Scientific, Waltham, MA, USA) and the primers are presented in Table S1B. Relative quantification of the copy number and the gene expression was performed with the 2^-ΔΔCt^ method using the *M. lusitanicus* actin gene (CBS277.49v2.0 genome database: scaffold_07: 2052804-2054242) as a reference (60). Amplification conditions involved 95 °C for 3 min followed by 40 cycles at 95 °C for 15 sec, 60 °C for 30 sec and 72 °C for 30 sec. Melting curve analysis was performed at 65 to 95 °C, with a 0.5 °C increment. All experiments were performed in biological and technical triplicates.

### Knock-out of the *cotH* genes using the CRISPR-Cas9 method

Knock-out mutants were constructed by exchanging partially the coding regions of the *cotH* genes to a functional *pyrG* gene (CBS277.49v2.0 genome database ID: Mucci1.e_gw1.3.865.1), which complements the uracil auxotrophy of the applied strain. This gene replacement was carried out by the homology-driven repair (HDR) following a CRISPR-Cas9 strategy described previously (29, 61). Protospacer sequences designed to target the DNA cleavage in the *cotH1*, *cotH2*, *cotH3*, *cotH4* and *cotH5* genes are presented in Table S2A. Using these sequences, Alt-R CRISPR crRNA and Alt-R CRISPR-Cas9 tracrRNA molecules were designed and purchased from Integrated DNA Technologies (IDT, Coralville, IA, USA). To form the crRNA:tracrRNA duplexes (i.e. the gRNAs), the Nuclease-Free Duplex Buffer (IDT, Coralville, IA, USA) was used according to the instructions of the manufacturer. Deletion cassettes functioning also as the template DNAs for the HDR were constructed by PCR using the Phusion Flash High-Fidelity PCR Master Mix (Thermo Fischer Scientific, Waltham, MA, USA). First, two fragments being upstream and downstream from the protospacer sequence of the corresponding *cotH* gene and the entire *pyrG* gene along with its promoter and terminator sequences were amplified using the primers listed in Table S1B. The amplified fragments were fused in a subsequent PCR using nested primers (see Table S1B in the supplemental material) where the ratio of concentrations of the fragments was 1:1:1. For each transformation procedures, 5 μg template DNA, 10 μM gRNA and 10 μM Cas9 nuclease enzyme (Alt-R S.p. Cas9 Nuclease, IDT, Coralville, IA, USA) were introduced together into the *M. lusitanicus* MS12 strain by PEG-mediated protoplast transformation (29, 62). Potential mutant colonies were selected on solid YNB medium by complementing the uracil auxotrophy of the MS12 strain. From each primary transformant, monosporangial colonies were formed under selective conditions. Disruption of the *cotH* genes and the presence of the integrated *pyrG* gene were proven by PCR using the primers listed in Table S1B and sequencing of the amplified fragment. Sequencing and PCR revealed that the CRISPR-Cas9-mediated HDR caused the expected modification (i.e., disruption of the *cotH* genes by the integration of the *pyrG*) in the targeted sites. Real-time quantitative reverse transcription PCR (qRT-PCR) analysis indicated the lack of *cotH* transcripts in all transformants.

### Complementation

To complement the knock-out of the *cotH3* and the *cotH4* genes, autonomously replicating vectors were used, where the complementing gene constructs were maintained episomally. *CotH3* and *cotH4* with their promoter and terminator sequences were amplified by PCR using the Phusion Flash High-Fidelity PCR Master Mix (Thermo Scientific) and the primer pairs *McCotH3_compl_fw* and *McCotH3_compl_rev* and *McCotH4_compl_fw* and *McCotH4_compl_rev*, respectively, and were ligated separately into pJet1.2 cloning vectors (Thermo Scientific) resulting the constructions pJet1.2-McCotH3 and pJet1.2-McCotH4, respectively. We further ensured that a unique rare-cutting restriction site is present in both plasmids (*Not*I) which should facilitate easy insertion of complementing genes. We have therefore combined in a single vector the auxotrophic selection marker *leuA* allowing primary selection of numerous transformants with our gene of interest resulting in plasmids pMCcotH3leuAcomp and pMCcotH4leuAcomp. The plasmid construction was introduced to the MS12-Δ*cotH3*+*pyrG* and MS12-Δ*cotH4*+*pyrG* disruption strains by PEG-protoplast transformation. In experiments with a plasmid for complementation of *cotH3* and *cotH4* gene deletion 3 μg DNA was added to the protoplasts in a transformation reaction. Transformants were transferred to minimal media (YNB) considering the phenomenon of complementation of auxotrophy after successful transformation and confirmed by PCR and qPCR. The selection method is based on the complementation of the leucine auxotrophy. The created knock-out and complemented *M. lusitanicus* strains are listed in Table S2B.

### Electron microscopy and quantitative analysis of the wall thickness

Pellets from isolated spores were immersed into a 2% paraformaldehyde (Sigma, St. Louis, MO, USA) and 2.5% glutaraldehyde (Polysciences, Warrington, PA, USA) containing modified Karnovsky fixative in phosphate buffer. The pH of the solution was adjusted to 7.4. Samples were fixed overnight at 4 °C, then briefly rinsed in distilled water (pH7.4) for 10 minutes and fixed in 2% osmium tetroxide (Sigma Aldrich, St. Louis, MO, USA) in distilled water (pH7.4) for 60 minutes. After osmification, samples were rinsed in distilled water for 10 minutes again, then dehydrated using a graded series of ethanol (Molar, Halasztelek, Hungary) from 50% to 100% for 10 minutes in each. Afterwards, all spore pellets were proceeded through in propylene oxide (Molar, Halasztelek, Hungary), then embedded in an epoxy-based resin, Durcupan ACM (Sigma Aldrich, St. Louis, MO, USA). After polymerization for 48 hours at 56 °C, resin blocks were etched, and 50 nm thick ultrathin sections were cut on an Ultracut UCT ultramicrotome (Leica, Wetzlar, Germany). Sections were mounted on a single-hole, formvar-coated copper grid (Electron Microscopy Sciences, Hatfield, PA, USA). For a better signal-to-noise ratio, 2% uranyl acetate (in 50% ethanol (Molar, Halasztelek, Hungary); Electron Microscopy Sciences) and 2% lead citrate (in distilled water; Electron Microscopy Sciences, Hatfield, PA, USA) were used. Ultrathin sections from the pellets were screened at 1.000–3.000× magnification on a JEM-1400Flash transmission electron microscope (JEOL, Tokyo, Japan) until 70 individual spore cross-sections were identified from each sample. For quantitative measurements of the major and minor axes, area, circularity 1.000×−3.000× magnification images were used. For the thickness measurements of the different layers of the spore wall (Fig. 5D), images were recorded at 10.000× magnification using a 2k×2k Matataki (JEOL, Tokyo, Japan) scientific complementary metal-oxide-semiconductor camera. All the quantitative data were determined using the built-in measurements module of the TEM Center software (JEOL, Tokyo, Japan).

### Fluorescence staining

Four days old fungal spores were collected from MEA and washed three times with 1x sterile phosphate buffer saline (PBS; 137 mM NaCl, 2.7 mM KCl, 10 mM Na_2_HPO_4_, 2 mM KH_2_PO_4_, pH7.4) and were collected by centrifuging for 10 min at 4 °C 2000×g, 5 min. The pellet was resuspended in 0.5 ml of 1% (w/v) bovine serum albumin (FBS; Gibco) solution and incubated for 30 minutes at room temperature with constant rotation. Fungal spores (10^7^/ml) were washed three times with 1x PBS, then stained for 45 min in PBS containing 5 μg/ml CFW solution or 100 μg/ml ConA-FITC solution. After staining, samples were washed five times with PBS. For analysis, collected samples were centrifuged with 2000×g for 15 min and resuspended in 200 μl PBS supplemented with 0.05% Tween-20 (Reanal, Budapest, Hungary). Fluorescence images of the stained fungal cells were taken with a Zeiss Axioscope 40 microscope and an Axiocam Mrc camera. A 350 nm excitation and 432 nm emission filters were used to detect chitin, while a 495 nm excitation and 515 nm emission filters were used after Con-A-fluorescein isothiocyanate staining. Samples were also measured with a FlowSight Imaging Flow Cytometer (Amnis^®^ ImageStream^®^X Mk II Imaging Flow Cytometer; Austin, Texas, US) and the associated IDEAS 6.2 software (63) was used for evaluation. For samples prepared by fluorescence microscopy, the intensity of the dyes for the different strains was determined using ImageJ2 (Fiji) (64). The mean fluorescence intensity of CFW was determined by microscopy. Significance was determined by unpaired t test.

### Interactions of *M. lusitanicus* with macrophages

The murine macrophage cell line J774.2 was cultivated in Dulbecco’s minimal essential medium (DMEM, Lonza, Basel, Switzerland) supplemented with 10% heat-inactivated fetal bovine serum (FBS, Biosera, Kansas City, MO, USA) and 1% penicillin/streptomycin solution (Sigma-Aldrich, St. Louis, MO, USA) at 37 °C, 5% CO_2_ and 100% relative humidity. The same medium was used in all the interaction experiments. Four hours before the experiment, J774.2 cells (2×10^5^ cells/ml) were freshly harvested and stained with CellMask Deep Red Plasma Membrane stain (Thermo Fischer Scientific, Waltham, MA, USA) following the instructions of the manufacturer, then seated on a 24-well plate. Fungal spores were freshly collected from 1-week-old MEA cultures and stained with Alexa Fluor 488 carboxylic acid, succinimidyl ester (Invitrogen, Waltham, MA, USA). Labelled macrophages and spores were co-incubated at a multiplicity of infection (MOI) of 5:1 for 90 min at 37 °C and 5% CO_2_. For analysis, collected samples were centrifuged with 1000×g for 10 min, then resuspended in 200 μl PBS supplemented with 0.05% Tween-20. Interaction and phagocytosis were measured using a FlowSight Imaging Flow Cytometer (Amnis^®^ ImageStream^®^X Mk II Imaging Flow Cytometer; Austin, Texas, US) and evaluated with the IDEAS Software (Amnis^®^ ImageStream^®^X Mk II Imaging Flow Cytometer; Austin, Texas, US). Data from 10.000 events per sample were collected and analyzed. The number of engulfed cells was determined by examining 200 hundred pictures of individual macrophages, while the phagocytic index (PI) was determined using the following formula: PI = [mean spore count per phagocytosing cell] × [% of phagocytosing cells containing at least one fungal spore].

To analyze the phagosome acidification of J774.2 cells by flow cytometry, fungal spores were labelled with pHrodo Red succinimidyl ester (Invitrogen, Waltham, MA, USA), according to the manufacturer’s instructions. CellMask Deep Red Plasma Membrane stain (Thermo Fischer Scientific, Waltham, MA, USA) labelled macrophages and FITC labelled spores were co-incubated at a MOI of 5:1, for 120 min at 37 °C and 5% CO_2_. The ratio of the phagocytosing cells was also determined at 120 min under the same conditions of interactions, except that fungal spores were stained with CFW. Phagosome acidification was calculated as follows: [% of pHrodo^™^ Red positive cells] / [% of phagocytosing cells] × 100.

To assay the survival of the fungal spores, the interaction of J774.2 cells and the spores was performed using a MOI of 5:1 for 180 min. After the interaction, cells and spores were collected and macrophages were lysed with sterile distilled water. Serial dilutions were prepared from the spore suspensions and plated on MEA to quantify the colony-forming units (CFU). Survival of spores was calculated using the following formula: Survival = CFU_interaction_ × 100 / CFU_control_, where CFU_interaction_ is the CFU of samples co-incubated with macrophages, while CFU_control_ is the CFU of control samples, incubated under the same conditions but without macrophages.

### Survival assay in *Drosophila melanogaster*

*Drosophila* stocks were raised and kept following the infection on standard cornmeal agar medium at 28 °C. Spore suspensions were prepared in sterile PBS from 7-day-old cultures grown on YNB plates (supplemented with 0.5 g/L leucine, if required) at 28 °C. The *Drosophila* Oregon R strain, originally obtained from the Bloomington stock center, was used throughout the experiments. Infection was performed by dipping a thin needle in a suspension of fungal conidia (10^7^ conidia/mL) or PBS as the uninfected control, and subsequently, the thorax of the anaesthetized fly was collected. Flies were counted at a different time to monitor survival. Flies were moved into fresh vials every other day. Each experiment was performed with 60 flies. The results shown are representative of at least three independent experiments.

### Survival assay in *Galleria mellonella* larvae

Spores were resuspended in insect physiological saline (IPS; 50 mM NaCl, 5 mM KCl, 10 mM EDTA and 30 mM sodium citrate in 0.1 M Tris–HCl, pH6.9) (65). *G. mellonella* larvae (BioSystems Technology, TruLarv) were inoculated with 10^5^ fungal cells in 20 μl IPS via the last proleg using 29-gauge insulin needles (BD Micro-Fine). For each *M. lusitanicus* strain, 20 larvae were infected. For IPS-treated (uninfected) and witness control (no injections, uninfected), 20 animals were utilized too. Larvae were maintained at 28 °C, and their survival was monitored daily for 6 days. The results shown are representative of at least three independent experiments.

### *In vivo* virulence studies in diabetic ketoacidotic (DKA) mouse model

Male ICR (CD-1^®^) outbred mice (≥20 g) were all purchased from Envigo and housed in groups of 8 each. Mice were rendered DKA with a single intra peritoneal injection of 190 mg/kg streptozotocin in 0.2 ml citrate buffer (pH4.2) 10 days before the fungal challenge. Glycosuria and ketonuria were confirmed in all mice 7 days after streptozotocin treatment. Furthermore, on days −2 and +3 relative to infection, mice were given a dose of cortisone acetate (250 mg/kg s.c.). Mice were given 50 ppm enrofloxacin (Baytril; Bayer) added to the drinking water from Day −3 to Day 0 ad libitum. Ceftazidine antibiotics were added (5 mg/dose/0.2mL SQ) from Day 0 and continued until Day +8. DKA mice were infected intratracheally with a target inoculum of 2.5 × 10^6^ (1X10^8^ spores/ml) fresh spores in 25 μl PBS after sedation with isoflurane gas.

### Ethics statement

All animal studies were approved by the Institutional Animal Care and Use Committee (IACUC) of the Los Angeles Biomedical Research Institute at Harbor-UCLA Medical Center according to the NIH guidelines for animal housing and care (Project 11671).

### Statistical analysis

All measurements were performed in at least two technical and three biological replicates. Statistical significance was analyzed by t-tests or One-way ANOVA followed by Dunnett’s multiple comparisons test using Microsoft Excel of the Microsoft Office package or GraphPad Prism 7.00 (GraphPad Software, La Jolla, California USA) as appropriate. *P* values less than 0.05 were considered statistically significant. In *in vivo* survival experiments, differences between the pathogenicity of the fungal strains were compared by the Log Rank test. *P* values less than 0.05 were considered statistically significant.

## SUPPLEMENTAL MATERIAL

Supplemental material is available online only.

**FIG S1**, TIFF file, 2.62 MB.

**FIG S2**, TIF file, 2.57 MB.

**FIG S3**, TIF file, 2.26 MB.

**FIG S4**, TIF file, 2.46 MB.

**FIG S5**, TIF file, 442 KB.

**FIG S6**, TIF file, 2.92 MB.

**FIG S7**, TIF file, 2.39 MB.

**TABLE S1**, excel file, 14 KB

**TABLE S2**, excel file, 12 KB

**FILE S1**, excel file, 44.6 KB

All relevant data are within the manuscript and its Supporting Information files.

## ACKNOWLEDGMENT

This study was supported by the grants NKFIA K131796, ELKH 2001007 and ITM NKFIA TKP-2021-EGA-28 to T.P. and by the Public Health Service grant R01 AI063503 to A.S.I. Publication was supported by the University of Szeged Open Access Fund (grant no. 5752). GN is grateful for the support of the Premium Postdoctoral Fellowship Program of the Hungarian Academy of Sciences (460050). C.S. is supported by the ÚNKP-20-4-I and ÚNKP-22-4-SZTE-523 New National Excellence Program of the Ministry for Innovation and Technology from the source of the National Research, Development and Innovation Fund. R.P. and K.S. were supported by the “National Talent Programme” with the financial aid of the Ministry of Human Capacities (NTP-NFTÖ-22-B-0027 and NTP-NFTÖ-22-B-0069).

We would like to thank Szilárd Szebenyi for his help in the performance of the figures.

The authors are grateful to Victoriano Garre and DoE JGI for contributing the unpublished data of the *Mucor lusitanicus* CBS277.49 v3.0 genome used in the phylogenetic analysis of the *cotH* sequences. Sequence data were produced by the US Department of Energy Joint Genome Institute https://www.jgi.doe.gov/ in collaboration with the user community. The work conducted by the U.S. Department of Energy Joint Genome Institute, a DOE Office of Science User Facility, is supported by the Office of Science of the U.S. Department of Energy under Contract No. DE-AC02-05CH11231.

A.S.I. owns shares in Vitalex Biosciences, a start-up company that is developing immunotherapies and diagnostics for mucormycosis. The remaining authors declare no competing interests.

Resources were provided by T.P., G.N., R.P., R.S., U.B., A.S.I. Conceptualization, methodology and supervision were performed by T.P., G.N., A.S.I., C.V. Validation of the results, writing, supervising, and editing the manuscript were carried out by T.P., C.S., G.N., R.P., R.S., A.S.I., C.V. Investigation was performed by C.S, S.K-V., O.J., Y.G., T.G., K.S., M.H., U.B. Figures and tables were arranged by C.S., S.K., R.P., A.S.I. All authors collected data, provided resources, participated in data curation, formal analysis and/or contributed to the revision of the manuscript.

## Legends for Supplemental Materials

**FIG S1 Maximum Likelihood tree showing phylogenetic relationships of *cotH* gene sequences.** Only bootstrap values >95 % from maximum-likelihood analyses are shown above the branches.

**FIG S2 Transcription analysis of the *M. lusitanicus cotH* genes.** (A) Relative transcription level of *cotH1-5* genes on minimal medium at 28 °C during the first four day of cultivation and (B) RT-qPCR analysis of *cotH1-5* genes after treatment with human serum at the second cultivation day on minimal medium containing 10% human serum. (C) Relative transcription level of *cotH1-17* genes on minimal medium at 28 °C on the second day of cultivation. (D) Relative transcription level of *cotH1-17* genes on the first and second day of cultivation on minimal medium containing 1 and 2.5% glucose. Values presented are from three independent cultivations (error bars indicate standard deviation).

**FIG S3 Growth of *cotH* mutants under microaerophilic conditions.** (A) Growth test of the *cotH* mutants (MS12-Δ*cotH1*+*pyrG*, MS12-Δ*cotH2*+*pyrG*, MS12-Δ*cotH3*+*pyrG*, MS12-Δ*cotH4*+*pyrG*, respectively) and the control MS12+*pyrG* strain on minimal medium under microaerophilic condition. Plates were incubated for two days on solid media at 28 °C in continuous light in a GasPak Anaerobic System. (B) Colony diameters of the strains plotted in mm, after two days on incubation under microaerophilic condition and under aerobiosis, as well. Absolute growth of the strains in microaerophilic and aerophilic cultivation was calculated based on the colony diameter data shown here as follows: [the growth of the mutants under the given condition] / [the growth of the control under the given condition] × 100. 100% was taken to be the growth of the control under the given condition. (m. ae. = microaerophilic condition; ae. = aerophilic conditions). (C) Absolute growth of the MS12-Δ*cotH4*+*pyrG* compared to the control MS12+*pyrG* in both microaerophilic and aerobic conditions. Values indicated with asterisks significantly differed from the corresponding value of the MS12+*pyrG* strain according to two-sample, paired t-test, where ** *P* ≤ 0.01. Error bars indicate standard deviation.

**FIG S4 Changes in the spores after complementation of *cotH4* disruption.** A) Transmission electron microscopic pictures of spores after complementation of *cotH4* gene: A/1) MS12-Δ*cotH4*+*pyrG;* A/2) MS12-Δ*cotH4*+*cotH4*+*leuA;* A/3) MS12+*pyrG;* and changes in the cell wall thickness (B); Spore circularity (C); Cross-section (D); Longitudinal cross-section (E) and Profile area (F) of the spores after plasmid-based complementation of *cotH4* gene in *M. lusitanicus*. Error bars indicate standard deviation.

**FIG S5 Surface analysis of fungal spores with Calcofluor white (CFW).** Dye-emitted intensity of the *cotH4* mutant and MS12-Δ*cotH4*+*cotH4*+*leuA* spores after CFW staining compared to the control (MS12+*pyrG*). Values indicated with asterisks significantly differed from the corresponding value of the MS12+*pyrG* strain according to two-sample, paired t-test, where ** *P* ≤ 0.001. Error bars indicate standard deviation.

**FIG S6 *In vitro* interactions with macrophages.** (A) Average number of phagocytosed spores during *in vitro* interactions with macrophages. (B) Phagocytosis of *M. lusitanicus* spores by J774.2 macrophages. J774.2 cells and spores were stained with CellMask Deep Red Plasma Membrane Stain and Alexa Fluor 488 carboxylic acid, succinimidyl ester, respectively. Macrophages were identified by detecting fluorescence intensity on channel 11 (Intensity_MC_CH_11) while channel 2 (Intensity_MC_CH_2) was used to detect the spores. Cells and spores were co-incubated for 3 h. Fluorescent micrographs showing spores and J774.2 cells alone (red border) and in interaction (i.e., phagocytosis or attachment) (green border) were recorded during the imaging flow cytometry. (C) Phagocytic index values. (D) Survival of fungal spores co-cultured for 3 h with murine J774.2 cells. Error bars indicate standard deviation.

**FIG S7 Acidification of phagosomes during interaction with *cotH* mutant spores.** (A) J774.2 cells were co-incubated with the labelled spores and then the ratio of pHrodo^™^ Red + macrophages were examined by imaging flow cytometry. Phagosome acidification was calculated as follows: [% of pHrodo^™^ Red positive cells] / [% of phagocytosing cells] × 100. (B) Phagosomal acidification of macrophages after *in vitro* interaction with *cotH* mutant spores. Error bars indicate standard deviation.

**TABLE S1** (A) Characteristics and efficiency of the CRISPR-Cas9 mediated genome editing in the creation of *cotH* mutant strains. (B) Primers used in the present study.

**TABLE S2** (A) Sequences of the protospacers designed in the study and positions of the corresponding regions in the *Mucor lusitanicus cotH* genes. (B) *Mucor lusitanicus* strains used in the study.

^1^ Numbers present nucleotide positions downstream from the start codon of the *M. lusitanicus cotH1* gene (scaffold_5:3666732-3669040).

^2^ Numbers present nucleotide positions downstream from the start codon of the *M. lusitanicus cotH2* gene (scaffold_1:3072475-3074523).

^3^ Numbers present nucleotide positions downstream from the start codon of the *M. lusitanicus cotH3* gene (scaffold_3:4206378-4208599).

^4^ Numbers present nucleotide positions downstream from the start codon of the *M. lusitanicus cotH4* gene (scaffold_4:354982-357045).

^5^ Numbers present nucleotide positions downstream from the start codon of the *M. lusitanicus cotH5* gene (scaffold_4:2485275-2486992).

**FILE S1 Transmission electron microscopic analysis of the spores of the mutant strains.** Changes in the cell wall thickness of the *cotH3* and *cotH4* mutants by (transmission electron microscopic) TEM measurements. Spores of the mutants and the control strain (MS12+*pyrG*) were subjected to (TEM) image analysis.

